# Preparation of inexpensive, pre-stained molecular weight marker using proteins isolated from chicken egg for SDS PAGE and Western blot application

**DOI:** 10.1101/2024.07.30.604739

**Authors:** Karthika KB Balaji, Vini VJ Jain, Sundarraj SR Rangasamy, Gokila GR Rajendran, Vinothkumar VK Kittappa, Arumugam AM Muruganandam

**Affiliations:** MANIPAL SCHOOL OF LIFE SCIENCES; AFFIGENIX BIOSOLUTIONS CHENNAI

**Keywords:** Pre-stained protein marker, Protein ladder, Chicken egg, SDS-PAGE, Western blot

## Abstract

Molecular weight markers, which are utilized in protein chemistry applications, including gel filtration chromatography, SDS-PAGE, and Western blotting, determine the molecular weight of distinct proteins that have been separated. These markers act as standards and are made of a variety of differently sized proteins that lie within a wide range of sizes corresponding to known molecular weights. They are indispensable tools in techniques such as protein analysis and characterization, as they may be used to estimate the approximate molecular weight of unknown proteins by comparing their movement pattern on the gel to established standards. Molecular weight markers can be obtained from multiple sources. There are many ways to obtain molecular weight markers, such as recombinant expression from organisms such as *E. coli* or mammalian cells as well as purification of proteins from organisms such as *E. coli*, bovine plasma, porcine muscle, and bovine milk. Moreover, the chicken egg white and egg yolk consist of proteins of varying molecular weights. This study aims to employ cost effective natural products such as chicken egg to create easily developed, reasonably priced protein markers with molecular weights that range from 250 to 14 kDa.

---

Protein molecular weight markers, also known as protein standards or ladders, are tools which estimate the protein size in various laboratory technique such as gel filtration chromatography, Western blotting, and SDS-PAGE. These markers are used as standards to estimate the size of unknown proteins by comparing the migration pattern against it ^1^.

The protein markers currently being utilized for laboratory purposes are expensive however they are indispensable. Their primary function is to aid in comparison of band pattern of samples. In SDS-PAGE, both pre-stained and unstained protein markers serve as references to identify purified proteins. Pre-stained markers are typically labeled with a dye or stain beforehand, this allows them to be directly visualised without any additional staining required after electrophoresis. Remazol dyes, particularly the potassium salts of sulfato ethyl sulfone derivatives of anthraquinone sulfonic acid are used to stain proteins in gels by forming covalent linkage between the vinyl sulfone derivative of remazol dyes and protein functional groups under specific conditions. The covalent bond between the protein and dye are responsible for the high specificity in protein detection^2,3^.

Protein markers can be derived from recombinant DNA technology or from native biological sources, they are known as recombinant and natural markers respectively. Proteins of different sizes that have been produced and purified in recombinant heterologous hosts make up such as Beta-galactosidase, Phosphorylase B, Bovine Serum Albumin, Ovalbumin, Turkey Albumin, Carbonic Anhydrase, Soybean Trypsin Inhibitor, Lactalbumin, and Lysozyme. Protein markers made using recombinant proteins are often highly laborious and expensive, meanwhile natural markers isolate proteins from native sources, they are relatively inexpensive^4,5^.

Chicken eggs are known as an economical source of a variety of native found in both the egg white and yolk. The egg white contains approximately 54% Ovalbumin, 12% Ovatransferrin, and 3.5% each of Ovamucin and lysozyme^6^. The major component of egg yolk is Chicken IgY, an antibody isotype which provides passive immunity to the developing embryo. Chicken IgY has a molecular weight of 180 kDa, and is abundantly present (40-80mg per egg) in the egg yolk. Upon heat denaturation, IgY yields heavy chains of approximately 65-70 kDa and light chains of 22-30 kDa^7^. These proteins can be readily isolated in large quantities using simple chemical methods, combined in appropriate proportions, pre-stained, and can be employed as molecular markers in SDS PAGE and Western blot applications. This study describe the method of isolating diverse proteins of varying sizes from chicken eggs and characterized for intended use molecular weight markers in protein chemistry studies.

## Materials and methods

### Purification of IgY from Chicken Egg Yolk

The chicken egg yolk from which the protein was isolated was obtained from Gallus gallus domesticus, commonly known as Leghorn chicken. Upon carefully cracking open the eggs, a yolk separator was used to eliminate the egg white. The collected egg white was then preserved for subsequent isolation of egg white proteins. IgY was purified from the isolated egg yolk following a previously established protocol with minor adjustments^8^.

Briefly, excess egg white was blotted out using filter paper, and precise quantity of the egg yolk volume were recorded. Twice the volume of 1X PBS was added to the yolk and thoroughly mixed to ensure uniformity. The 3.5% PEG 6000 solution was added to the mixture and vortexed for two minutes. The mixture was then allowed to incubate at room temperature for 10 minutes with shaking at 200 rpm. The content was centrifuged at 3500 rpm for 15 minutes at 4°C to collect the supernatant in a clean tube.

The collected supernatant was treated with 8.5% PEG solution and the content were incubated and centrifuged as done in the previous steps. The supernatant was discarded and the pellet was resuspended in 10 ml of phosphate buffered saline. 12% PEG 6000 was added to the suspension and incubated and centrifuged as done previously. The pellet was resuspended in 10 ml of phosphate buffered saline and stored at -20°C.

### Isolation of egg white Proteins

The egg white was isolated from the egg yolk, and transferred to a 500 ml beaker. The white was mixed with distilled water to get a 100 ml total volume. With the magnetic stirrer in place, this mixture was mixed slowly for half an hour, minimising foaming. After adding the egg white suspension to a sterile centrifuge tube, the mixture was centrifuged for 20 minutes at 4°C and 3,000 rpm. The mixture was used as the starting material for the further purification of various egg white proteins.

The supernatant from the egg white suspension was employed in the purification process of ovalbumin. The required amount of ammonium sulphate was gradually added, based on the volume of the supernatant, until a final concentration of 27.5% was reached. This solution was stirred gently using a magnetic stir. The salts were allowed to completely dissolve and then left to incubate at room temperature for 15 minutes. Following this, proteins that had precipitated were centrifuged for five minutes at 3,000 rpm. The supernatant was then gently transferred to another tube and ammonium sulphate precipitation was performed using the right amount of salt to achieve a final concentration of 55%. The pellet containing ovalbumin, which salted out, was then resuspended in 1 ml of water.

A portion of the collected supernatant from the egg white suspension was further utilized for the sequential isolation of lysozyme and Ovatransferrin, following a previously described method with minor adjustments^6^. To outline the procedure briefly, the supernatant was combined with 0.5g per 10 ml of cationic exchange resin (Livchrom SP Agarose CL6B; BioArtha Labs, Bengaluru), and the mixture was stirred for 12 hours at 4°C. Following the overnight incubation, the mixture underwent centrifugation at 3500 rpm for 20 minutes at 4°C. The resulting supernatant was collected and stored at 4°C for subsequent separation of ovotransferrin. The following step was the wash with 0.1M NaOH-Glycine buffer, pH 9.3. To extract the lysozyme from the cation exchange resin, 10 millilitres of 0.1M Glycine-NaOH buffer containing 0.5M NaCl was eventually utilised. Dialysis was then used to buffer-exchange the eluted solution with 1X PBS.

The lysozyme-free supernatant collected and stored in the previous step was pH adjusted to 4.75 using 3N HCl. After that, the content was centrifuged at 3,500 rpm for 30 minutes at 4°C. The supernatant was collected and was added with 5% ammonium sulfate and 2.5% citric acid and left overnight at 4°C to precipitate ovotransferrin. The contents were centrifuged at 3,500 rpm for 20 minutes at 4°C and 4 volumes of distilled water was added to the pellet. The pellet was buffer exchanged with 1X PBS by dialysis.

### Sodium Dodecyl Sulfate –Polyacrylamide Gel Electrophoresis (SDS-PAGE)

SDS-PAGE was conducted following the method outlined by Laemmli^9^. In brief, 10% resolving gel and 5% stacking gel were prepared and casted within glass plates. A comb was positioned in the stacking gel prior to polymerization to form wells. The gel-containing glass plates were placed within an electrophoresis device that was loaded with SDS running buffer. Following a suitable volume of 5X protein loading dye (250 mM Tris HCl, pH 6.8, 10% SDS, 50% Glycerol, 0.5% Bromophenol Blue), protein samples were put into the corresponding wells.

After being supplied by electric power for one and a half hours, the electrophoresis device was used to run the dye through the resolving gel. Post-electrophoresis, the gel was removed from the glass plates, and the stacking gel was carefully excised.

### Silver Staining of SDS PAGE Gel

As part of the silver staining technique, the gel was immersed in a fixing solution (made up of 40% ethanol, 10% acetic acid, and 50% deionized water) at room temperature and gently agitated for 20 minutes. following this the gel is transferred to a newly made sensitizing solution (made up of 6.8 g sodium acetate, 30 ml ethanol, 0.2 g sodium thiosulphate, and 500µL glutaraldehyde in 100 mL distilled water) and gently shaking it for 20 minutes at room temperature. The gel was then allowed to soak in a staining solution (0.25% silver nitrate) for 20 minutes after being rinsed twice with distilled water. The gel was placed in a newly made developing solution (consisting of 2.48 g sodium carbonate and 80 µL formaldehyde in 100 ml distilled water) after two more distilled water washes. The gel was placed in a stop solution (5% acetic acid) to cease the development process when it had developed to the appropriate contrast.

### Pre-staining of proteins with Remazol dye

To facilitate the identification and tracking of proteins during electrophoresis, marker proteins were pre-stained using Remazol dyes following the method described earlier with minor modification^10^. 5-10 mg of individual protein standards were dissolved in 250 microliters of a 100 mM sodium carbonate solution (pH adjusted to alkaline). After that, this solution was combined with 50μL of the dye, and it was incubated for 30 minutes at 60°C.

Following this, the reaction was halted by adding 10 mg of crystalline lysine and further incubating the mixture for 10 minutes at 60°C. Once each protein marker was prepared, 1-2μL of each reaction mixture was combined. These combined markers were then reconstituted in 15μL of SDS sampler buffer, achieving a concentration of 2-4μg of protein standard per sample. The markers were subsequently analyzed using SDS-PAGE.

### Transfer of proteins from SDS-PAGE to western blot membrane

The SDS-PAGE was used to run the prestained protein markers before they were transferred onto PVDF or nitrocellulose membranes (BioRad) at 80 mA for one hour using the transfer buffer Tris-glycine SDS (48 mM Tris, 39 mM glycine, 0.037% SDS) with 20% methanol. After transfer step, the membrane was simply visualized to see the transfer of all the four bands from the gel to the membrane.

## Results and discussion

This study attempted to isolate chicken egg proteins in order to use them as molecular weight markers. Two separate components of the chicken egg were utilized to isolate the desired proteins: yellow yolk and egg white. At first, the yolk was separated from egg white and subsequently the IgY antibodies were isolated from yolk that yielded an average of 60 mg of IgY per egg. Similarly, the egg white proteins lysozyme, ovalbumin and ovotransferrin were separated and their average yield was calculated as 7.5 mg, 870 mg and 98.2 mg per egg, respectively.

The resultant molecular weights (MW) of the protein mixture, which included IgY, lysozyme, ovalbumin, and ovotransferrin, were ascertained by contrasting them with a commercial molecular marker. The isolated proteins were mixed with 5X non-reducing SDS dye and separated through SDS-PAGE gel. The resulting resolved gel yielded lysozyme migrated at 14 kDa, ovalbumin at 35 kDa, ovotransferrin at 65 kDa and chicken IgY at 250 kDa (Figure 1).

**Figure 1.**
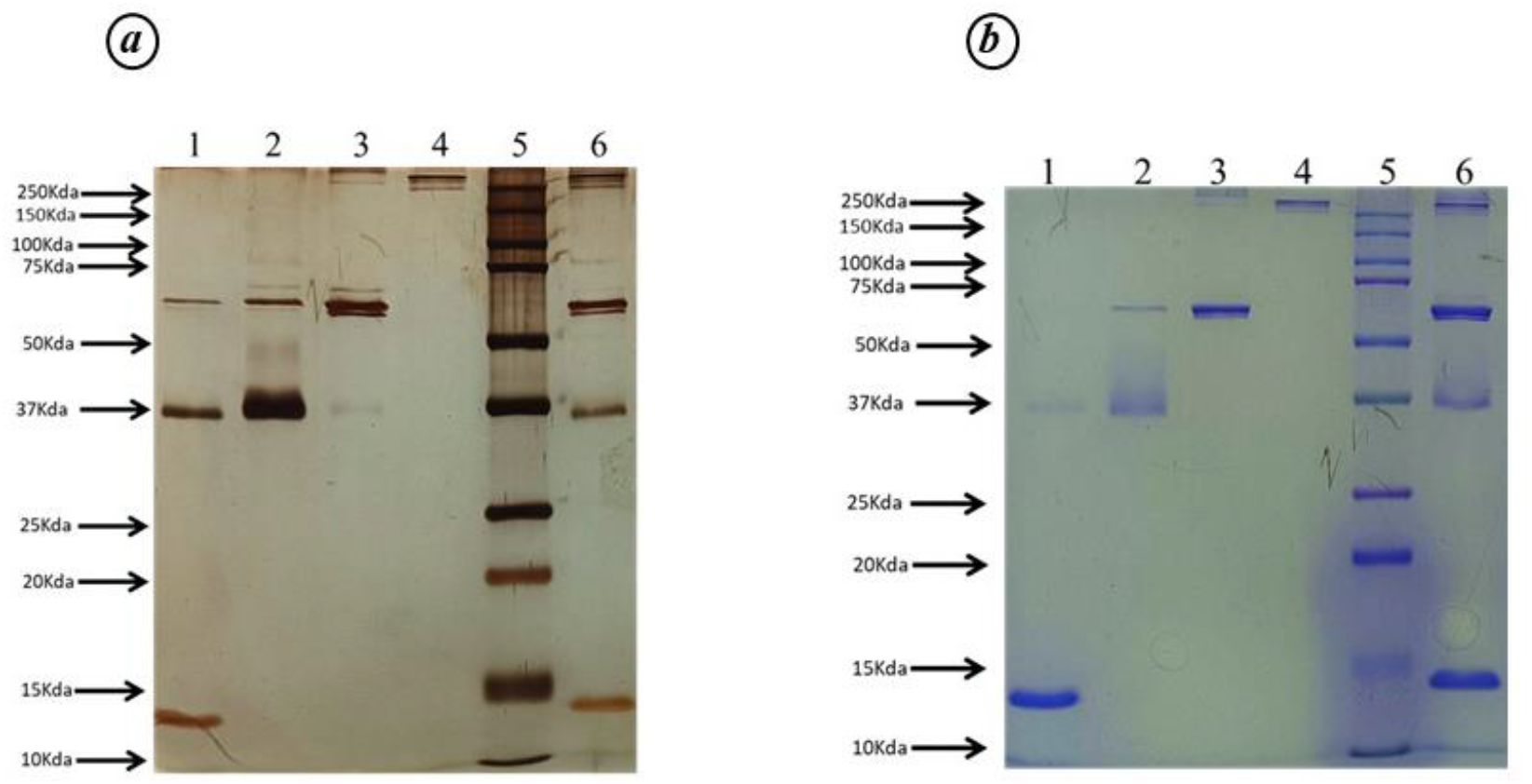
The SDS-PAGE (10%) analysis (a) Silver stained and (b) Coomasie stained, showing purified egg proteins lysozyme, ovoalbumin, ovotransfeITin and chicken IoY (Lanes 1-4 respectively), commercial protein marker (Lane 5) and marker prepared using the purified egg protein (Lane 6).

Throughout the isolation procedure, slight contamination was noted between lysozyme and ovalbumin, as well as between ovalbumin and ovotransferrin. Subsequent to protein isolation, an optimization step was undertaken to blend them in suitable concentrations, yielding a marker featuring four distinct bands visible in both Coomassie and silver-stained gels (Figure 1, Lane 6). Before application, the marker was pre-stained and observed via SDS-PAGE (Figure 2A), and its suitability for Western blot analysis was evaluated (Figure 2B). After pre-staining of the protein fraction of purified ovalbumin, two non-specific bands or artifacts (one around 16 kDa and another around 10 kDa) were showed up in SDS-PAGE. However those bands were not appeared in the transferred membrane.

**Figure 2.**
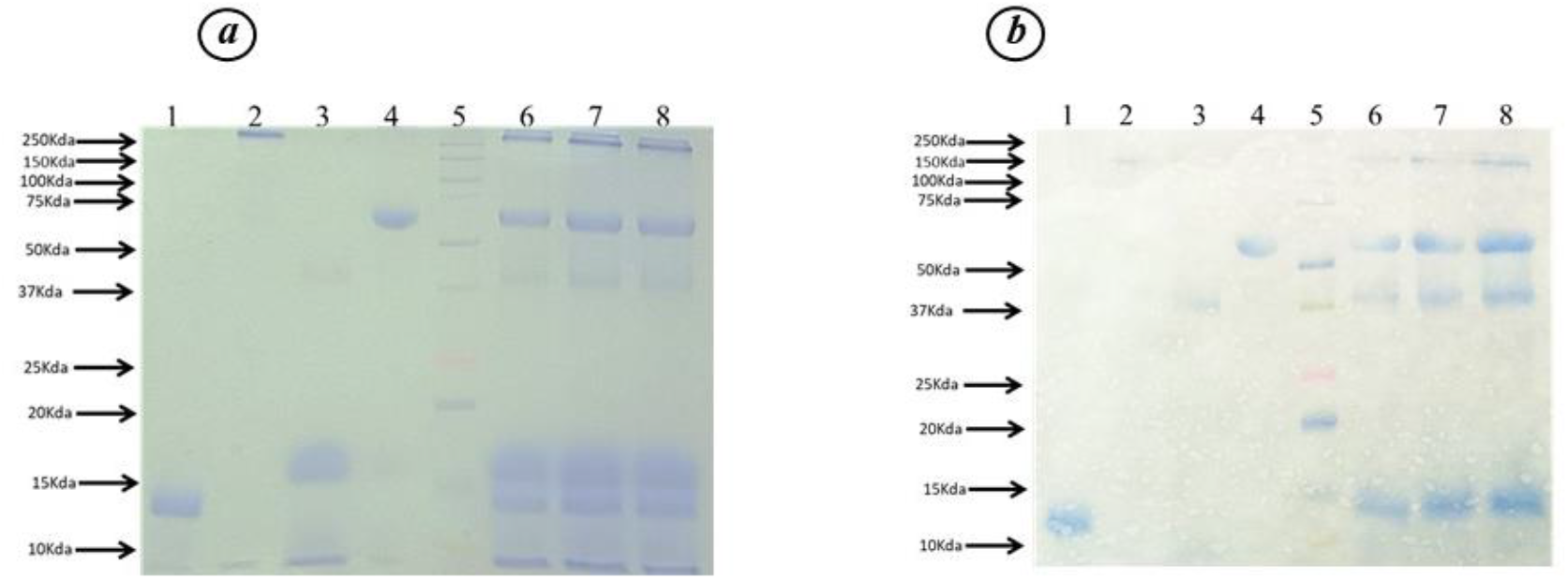
(a) The SDS-PAGE gel (10%) showing resolved pre-stained purified proteins lysozyme, chicken lg Y, ovalbllillin and ovotransfe1Tin in the lanes 1-4 respectively and the prepared pre-stained marker with total protein of 50, 60 and 80μg in the lanes 6-8 respectively. Commercial marker was loaded in the lane 5 (b) The c01Tesponding PVDF membrane showing the transfe1Ted pre-stained bands.

The study aimed to utilize the variety of proteins found in chicken eggs in order to produce and apply molecular weight protein markers for SDS-PAGE and Western blot procedures. The methodology involved the separation of proteins from the egg white and egg yolk sequentially. Lysozyme, ovalbumin, and ovotransferrin were isolated from the egg white, while IgY was isolated from the egg yolk using PEG6000. These extraction techniques were relatively easy to use and reasonably priced, making them appropriate for use even in smaller laboratory settings. There was some slight protein contamination in the lysozyme and ovalbumin fractions, but this was deemed negligible when the proteins were combined together to create the marker.

During gel electrophoresis, the protein marker, also known as the ladder, is used to approximate the size of unknown protein molecules. The ladder progresses through the gel and the purified proteins of varying sizes align based on their molecular weight. The larger the protein, the slower it migrates and it stays closest to the loading well. The SDS-PAGE buffer system that is used determines how the proteins migrate through the gel. Upon staining with dyes or silver nitrate, the protein bands serve as reference points to estimate the sample size.

The resulting marker displayed four distinct bands ranging from 15 kDa to 250 kDa, proving its value for various experiments. Pre-staining these markers contributed to their visualization during electrophoresis, eliminating the need for a separate staining step and simplifying the confirmation of protein marker transfer from the gel to the membrane in Western blotting procedures.

## Conclusion

In gel electrophoresis investigations, egg-derived protein markers might be a crucial instrument for quantifying protein size. These refined proteins are solid reference points for studies and electrophoretic transfers in western blot analysis due to their precisely determined molecular weights. In order to better understand protein functions and mechanisms and to understand protein functions and mechanisms, as well as in biochemistry and life sciences laboratories. This study shows the development of an easy to use, inexpensive, and pre-stained protein marker derived from chicken eggs, which is an easily accessible protein source. Its efficacy was proven by its successful use in Western blot and SDS-PAGE studies, indicating its potential use in a variety of research endeavors. This work also require improvement in terms of incorporating additional bands with molecular weights situated between the existing bands within the current specified range, as well as enhancing purity.

## Acknowledgement

Authors thank the management of Affigenix Biosolutions Pvt Ltd for funding this study.

## Conflict of Interest

All the participating Authors Declare no conflict of interest.

## Notes

### Competing Interest Statement

The authors have declared no competing interest.

